# Gene losses resulting in specialized metabolism in the earliest divergent symbiotic *Frankia* clade can be linked to its low saprotrophic capabilities

**DOI:** 10.1101/2022.05.25.493452

**Authors:** Fede Berckx, Thanh Van Nguyen, Rolf Hilker, Daniel Wibberg, Kai Battenberg, Jörn Kalinowski, Alison Berry, Katharina Pawlowski

## Abstract

*Frankia* cluster-2 are diazotrophs that engage in root nodule symbiosis with host plants of the Cucurbitales and the Rosales. They are rarely found in the soil in the absence of their hosts. Previous studies have shown that an assimilated nitrogen source, presumable arginine, is exported to the host in nodules of *Datisca glomerata* (Cucurbitales), but not in the nodules of *Ceanothus thyrsiflorus* (Rosales). To investigate if an assimilated nitrogen form is commonly exported by cluster-2 strains, and which nitrogen source would then be exported to *C. thyrsiflorus*, gene expression levels, metabolite profiles and enzyme activities were analysed.

We found that the export of assimilated nitrogen in symbiosis is a common feature for *Frankia* cluster-2 strains, but which source is host-plant dependent. We also identified several gene losses.

The ammonium assimilation via the GS/GOGAT cycle for export to the host, entails a high demand of 2-oxoglutarate from the TCA cycle. This specialised metabolism seems to have led to genome reduction: we show that *Frankia* cluster-2 strains have lost the glyoxylate shunt and succinate semialdehyde dehydrogenase, leading to a linearization of the TCA cycle. This could explain the low saprotrophic potential of *Frankia* cluster-2.

## Introduction

Root nodules are formed as the result of a symbiotic relationship between a nitrogen-fixing soil bacterium and its host plant. All root nodule-forming plants belong to a single clade, which encompasses the orders Fabales, Rosales, Fagales, and Cucurbitales (Griesman *et al.* 2018). While legumes (Fabaceae, Fabales) and *Parasponia* (Cannabaceae, Rosales) engage with rhizobia, the actinorhizal symbiosis is established between a diverse group of plants within the Rosales, Fagales and Cucurbitales, and their endosymbiont *Frankia.* The bacterial genus *Frankia* can be split into four phylogenetically distinct clades which are referred to as cluster-1 to −4 (Normand *et al.* 1996; Nguyen *et al.* 2016). The first three clusters represent symbiotic strains, which roughly coincide with their host specificity. Host plants of cluster-1 are solely found within actinorhizal Fagales: i.e. symbiotic members within the families Betulaceae, Casuarinaceae, and Myricaceae except for *Gymnostoma* spp. (Casuarinaceae) and *Morella* spp. (Myricaceae). The host range of *Frankia* cluster-2 is much broader. They engage in symbiosis with all actinorhizal Cucurbitales, i.e. the families Datiscaceae and Coriariaceae. Cluster-2 also engages with actinorhizal Rosales of the family Rosaceae and with *Ceanothus* spp. (Rhamnaceae). Most of the host plants of *Frankia* cluster-3 are found within the Rosales: symbiotic members of the families Elaeagnaceae and Rhamnaceae, except for *Ceanothus* spp. Cluster-3 can also engage in symbiosis with the two host plant genera in Fagales which do not engage with cluster-1 strains: *Gymnostoma* spp. and *Morella* spp. *Frankia* cluster-4 strains are not symbiotic (Normand *et al.*, 1996; Pozzi *et a*l., 2018). Aside from the broad host range, *Frankia* cluster-2 is particularly interesting because it represents the earliest divergent symbiotic clade (Sen *et al.*, 2014; Persson *et al.*, 2015; Gtari *et al.*, 2015; Nguyen *et al.*, 2016).

Recent phylogenomic studies support the hypothesis that the common ancestor to all nodulating plants was symbiotic, but the symbiosis was subsequently lost in the majority of the lineages (Griesmann *et al.*, 2018; van Velzen *et al.*, 2018). This begs the question: was the original symbiont *Frankia* or rhizobia? The polyphyly of the oxygen protection system for nitrogenase in nodules indicates that the original symbiont would have been *Frankia* (van Velzen *et al.*, 2019). The nitrogenase enzyme complex carries out nitrogen fixation. It is a high energy demanding process and the supply of the required ATP is best met by aerobic respiration (Shah and Brill, 1997). However, the enzyme complex is rapidly degraded under oxygen exposure, thus leading to the oxygen dilemma of nitrogen fixation. *Frankia* can provide oxygen protection for nitrogenase independently of the host plant, by the formation of vesicles, while rhizobia cannot. *Frankia* vesicles are surrounded by multilayered envelopes, which contain hopanoids (Berry *et al.*, 1993). The number of layers is correlated with the oxygen tension, indicating that they act as gas diffusion barriers (Parsons *et al.*, 1987).

Since in actinorhizal nodules both plant and endosymbiont can contribute to oxygen protection of nitrogenase, evolution has led to a wide variety of systems (Pawlowski and Demchenko, 2012), even among host plants of the same cluster. In *Frankia* cluster-2 host plants of the Rosales, like *Ceanothus integerrimus* (Rhamnaceae), the vesicles seem to play the main role in oxygen protection of nitrogenase. *Frankia* will form non-septate vesicles with multilaminate envelopes. They are pear-shaped and located at the periphery of the infected cells (Strand and Laetsch, 1977). In contrast, in host plants of the Cucurbitales, e.g. *Datisca glomerata* (Datiscaceae), both partners are involved in oxygen protection. The vesicles have a thin, single laminate envelope, and are arranged in a radial orientation around the central vacuole of the infected cell. A thick layer of mitochondria encloses the vesicle zone, thereby minimising the exposure of vesicles to the plant cytosol and providing physiological oxygen protection (Silvester *et al.*, 1999). Due to these structural differences, the oxygen protection mechanisms can cause oxidative and nitrosative stress, which requires detoxification mechanisms. In legume nodules, the production of reactive oxygen species (ROS) was enhanced by the symbiotic haemoglobins that are part of the oxygen protection system of nitrogenase (Günther *et al.*, 2007). *Frankia* produces bacterial truncated haemoglobin, *trHbO*, during symbiosis (Pawlowski *et al.* 2007). The homologue of *trHbO* in *Mycobacterium tuberculosis* has been proposed to act as an oxygen diffusion facilitator for oxidases under hypoxic conditions (Liu *et al.*, 2004; Pathania *et al.*, 2002).

During symbiosis, the host plant is provided with fixed nitrogen in exchange for photosynthates. In legumes and most actinorhizal plants, such as *Alnus glutinosa* (Betulaceae, Fagales), the fixed nitrogen is exported to the cytosol of infected cells as ammonium (Guan *et al.*, 1997; Alloisio *et al.*, 2010; Udvardi and Poole, 2013; Lui *et al.*, 2018; Hay *et al.*, 2020). In the cytosol, ammonium is assimilated and transported to the xylem in different forms depending on the plant species. In actinorhizal nodules of *A. glutinosa*, it was shown that ammonium is assimilated *via* the plant glutamine synthetase/glutamate synthase (GS/GOGAT) cycle. The assimilation by *Frankia* only occurs at low levels (Guan *et al.*, 1997, Alloisio *et al.*, 2010). In *D. glomerata* several studies indicate that an assimilated form of nitrogen, presumably arginine, is exported from the endosymbiont to the cytosol of the infected cell and then transported to the surrounding uninfected cells (Berry *et al.*, 2004; Berry *et al.*, 2011; Salgado *et al.*, 2018). In the cytosol of the uninfected cells, arginine is broken back down to ammonium and re-assimilated via the GS/GOGAT cycle. A similar accumulation of arginine could be found in nodules of *Coriaria myrtifolia* (Coriariaceae; Wheeler and Bond, 1970). In conclusion, the export of assimilated nitrogen, specifically arginine, from the bacterial symbiont to the plant seems to be a common feature for actinorhizal Cucurbitales. However, no evidence for the export of arginine by *Frankia* has been found in nodules of *Ceanothus thyrsiflorus* (Rhamnaceae, Rosaeles) based on plant gene expression levels (Salgado *et al.*, 2018). In fact, it has been previously shown that asparagine accumulates in nodules of *Ceanothus velutinus* (Wheeler and Bond, 1970). So while arginine can be excluded as the form of exported nitrogen by *Frankia* in *Ceanothus*, the export of assimilated nitrogen cannot be excluded as such. We wanted to identify the exported nitrogen metabolite to find out whether the specialised metabolism of exporting an assimilated form of nitrogen was unique to host plants from the Cucurbitales or to *Frankia* cluster-2 symbioses in general.

The carbon and nitrogen metabolism are directly linked by the tricarboxylic acid (TCA) cycle. This cycle can be closed via several distinct reactions. In the classic TCA cycle, 2-oxoglutarate (2-OG) is the substrate to produce succinyl-coenzyme A (succinyl-CoA) by the activity of 2-OG dehydrogenase. Succinyl-CoA synthase then converts succinyl-CoA to succinate. An alternative TCA has been described for some bacterial species (Tian *et al.*, 2005), including rhizobia (Green *et al.*, 2000) which is advantageous under reducing conditions as required during nitrogenfixation. Here, succinic semialdehyde (SSA) is produced from 2-OG by 2-OG decarboxylase. SSA is then converted to succinate via SSA dehydrogenase (SSA-DH). An alternative pathway to produce succinate from 2-OG is the gamma-aminobutyrate (GABA) shunt (Xiong *et al.*, 2014), which again requires the activity of SSA-DH. Lastly, the TCA cycle can be closed by the glyoxylate shunt, where the production of 2-OG would be avoided (Zhang *et al.*, 2015). If not used for maintaining the TCA cycle, 2-OG can be pulled out and used for the assimilation of ammonium in the GS/GOGAT cycle which leads to the production of glutamate. Glutamate can then be used as a substrate in the biosynthesis of glutamine, asparagine, or arginine. In all root nodule symbioses examined thus far, TCA cycle intermediates are supplied to the microsymbiont by the host plant (Jeong *et al.*, 2004, Udvardi and Poole, 2013). The export of an assimilated form of nitrogen by *Frankia* cluster-2, such as arginine, would require more carbon skeleton input from the host plant than in systems where ammonium is exported directly, such as in *Frankia* cluster-1 symbiosis (Guan *et al.*, 1997). Most, but not all, of these carbon skeletons would be returned by the endosymbiont during the export of assimilated nitrogen. Thus, the export of assimilated nitrogen by the endosymbiont is not energy efficient for the symbiosis as a whole, as it requires more complex transport processes.

A better understanding of the *Frankia* cluster-2 symbiosis could improve the understanding of root nodule symbiosis as a whole. Unfortunately, the majority of *Frankia* cluster-2 strains cannot be cultured to this date, except for two strains: *Frankia coriariae* BMG5.1 (Gtari *et al.*, 2015) and *F. coriariae* BMG5.30 (Gueddou *et al.*, 2019). This implies that most analyses, such as gene expression studies via reverse transcription-quantitative polymerase chain reaction (RT-qPCR), must be conducted *in planta*.

This study aims to compare features of the nitrogen and carbon metabolism in nodules of *Frankia* cluster-2 host plants from two different orders: *D. glomerata* representing Cucurbitales, and *C. thyrsiflorus* representing Rosales. The former was nodulated by *Candidatus* Frankia californiensis Dg2, while the latter was nodulated by the closely related strain *Candidatus* F. californiensis Cv1 (Nguyen *et al.*, 2016; Normand *et al.*, 2017; Nguyen *et al.*, 2019). As a comparative system, nodules of *A. glutinosa* induced by *Frankia alni* ACN14a (Normand *et al.*, 2007) were included in some of the analyses. For this symbiosis, it is known that ammonium is exported from the bacterium to the host plant (Guan *et al.*, 1997). This study presents analyses of gene expression levels and of enzyme activities related to carbon and nitrogen metabolism, as well as protein modelling, which were performed to elucidate the metabolite exchange between host and symbiont.

## Material and Methods

### Biological material

*Ceanothus thyrsiflorus* and *Datisca glomerata* plants were grown as previously described (Salgado *et al.*, 2018). In brief, *C. thyrsiflorus* plant cuttings were obtained from a local nursery, Cornflower Farms (Elk Grove, CA, USA). The inoculum used, *Candidatus* Frankia californiensis Cv1, originated from nodules of *Ceanothus velutinus* plants collected in Sagehen Experimental Forest (Truckee, CA, USA; Nguyen *et al.*, 2019). Crushed nodules of *C. thyrsiflorus* propagated in the greenhouse were used to inoculate the new plants. *D. glomerata* plants were germinated from seeds and grown on nitrogen-poor soil (S-Jord, Hasselfors Garden AB, Hasselfors, Sweden). Eight weeks after germination, they were transferred to a 1:1 (v/v) soil/sand (1.2-2 mm Quartz; Rådasand AB, Lidköping, Sweden) mixture and inoculated with *Candidatus* F. californiensis Dg2 (Nguyen *et al.*, 2016) from crushed mature *D. glomerata* nodules. *Alnus glutinosa* seeds were obtained from Svenska Skogsplantor (Bålsta, Sweden). Plants were germinated from seeds, which were vernalized for two weeks, on nitrogen-poor soil. Six-week-old plants were inoculated with *Frankia alni* ACN14a (Normand *et al.*, 2007), which had been grown in basic propionate (BAP) medium without nitrogen (Benoist *et al.*, 1992) at 28°C for four weeks. Cells were spun down at 4,500 x *g* for ten minutes and washed twice with sterile milliQ H2O before use. All plants were grown at a light/dark rhythm of 16 h light/8 h dark, with 26°C during the light and 19°C during the dark phase. The greenhouse was kept at 60% relative humidity. For all plants, the nodulation status was confirmed before inoculation to ensure that no plant was already infected. All plants were watered with deionized water twice per week, with one-quarter strength Hoagland’s medium with 1 mM KNO3 once per week before nodulation, and with one-quarter strength Hoagland’s medium without nitrogen once per week after nodulation (Hoagland and Arnon, 1938). *Frankia coriariae* BMG5.1 (Gtari *et al.* 2015) was obtained from Deutsche Sammlung von Mikroorganismen und Zellkulturen (DSMZ) and maintained in a modified BAP medium (pH adjusted to 9) as recommended by the DSMZ (Supporting Information Table S1).

### RNA isolation, cDNA synthesis, and reverse transcription-quantitative polymerase chain reaction (RT-qPCR) analysis

Mature nodules were collected, flash-frozen in liquid nitrogen and stored at −80°C until RNA isolation. Nodules were ground in liquid nitrogen with mortar and pestle. RNA was isolated using the Spectrum Plant Total RNA kit (Sigma-Aldrich, Stockholm, Sweden) according to the manufacturer’s instructions except for an additional sonication step as described by Nguyen *et al.* (2019). Genomic DNA was removed using the On-Column DNase I digestion kit (Sigma-Aldrich, Sweden). cDNA was synthesised using the TATAA GrandScript cDNA Synthesis Kit according to the instructions of the manufacturer (TATAA Biocenter, Sweden), except for the RT-negative control in which the enzyme was omitted. RT-qPCR reactions were performed using Maxima SYBR Green/ROX (ThermoFisher Scientific) as previously described (Nguyen *et al.*, 2019): initial denaturation at 95°C for 10 min, followed by 40 cycles of 15s at 95°C, 30s at 60°C, and 30s at 72°C. The melt curve program was 15s at 95°C, 15s at 60°C, and 15s at 95°C. All reactions were performed with four technical replicates and three biological replicates. Primers were designed for sequences conserved between Dg2 and Cv1, while separate primers were designed for ACN14a. Primer efficiency between 95 and 105% was confirmed in all samples. All primer sequences used in this study are given in Supporting Information Table S2. The housekeeping gene *infC*, encoding the translation initiation factor IF-3 (Alloisio *et al.*, 20210), and the nitrogenase gene *nifD* were used for normalising the relative expression data, expressed as delta Ct in calculations and figures. Statistical analysis and visualization (Student’s T-test when comparing Cv1 against Dg2, or one-way ANOVA when comparing ACN14a, Cv1 and Dg2 followed by Tukey posthoc analysis) of the data were performed through RStudio using the packages dplyr and ggplot2 (Wickham, 2016; RStudio Team, 2020).

### Amino acid profiling

Samples were taken from *C. thyrsiflorus* plants to profile the total amino acid content of nodules, inoculated roots, and uninoculated roots. Samples were treated as described by Persson *et al.* (2016) and extracted as described by Hacham *et al.* (2002). Profiling analysis was performed at the Molecular Structural Facility, University of California, Davis (CA, USA), on a Hitachi L-8900 Amino Acid Analyzer (Ibaraki, Japan). For statistical analysis oneway ANOVA was performed on three biological replicates, followed by a Tukey posthoc analysis. Data were analysed and visualised in RStudio.

### Protein extraction, enzyme assays, and protein modelling and phylogeny

Liquid cultures of *F. alni* ACN14a and *F. coriariae* BMG5.1 were collected via centrifugation (4,500 x *g* for ten minutes) and washed thrice in sterile milliQ. Both culture pellets and flash-frozen nodule samples were ground in liquid nitrogen with mortar and pestle. 500 mg of each sample were transferred to 2 mL tubes and 500 μL of protein isolation buffer was added (100 mM MES, 15% (v:v) ethyleneglycol, 2% (v:v) 2-mercaptoethanol, 1 mM phenylmethylsulfonylfluoride (PMSF) (adapted from Berry *et al.*, 2004). Bacterial proteins were isolated using the ultrasonic homogenizer Sonoplus HD 2070 (Bandelin Electronic, Berlin, Germany) at 30% pulsing 2 times with 60s each time, on ice. Samples were centrifuged briefly to pellet cellular debris, and total soluble protein concentration was determined using bovine serum albumin (BSA) as standard (Bradford, 1976), and diluted to the same concentration.

Enzyme activity measurements for the transferase activity of glutamine synthetase (GS) were performed as adapted from Berry *et al.* (2004) and Romanov *et al.* (1998). Five μL of nodule extract, which corresponded to 100ng of soluble protein, were added to 100μL of the reaction mixture (20mM Tris-Acetate, 8.75mM Hydroxylamine, 1mM EDTA, 2.25mM MnCl_2_, 17.5mM NaH_2_AsO_4_, 2.75mM ADP, 35mM glutamine, pH 6.4) and incubated at 30°C. The reaction was stopped by adding 100μL of ferric reagent (3.2% w/v FeCl_3_, 4% w/v TCA, 0.5N HCl), after 1, 15, 30, 60, and 120 minutes, respectively. The negative control either did not contain the substrate, or contained nodule extract that had been boiled for 10 minutes at 95°C to completely denature the proteins. To distinguish between heat-labile plant GS activity or bacterial GSII activity, and heat-stable bacterial GSI activity, samples were incubated at 40°C, 50°C, or 60°C for 10 minutes as the treatments in the literature vary (Behrmann *et al.*, 1990; Edmands *et al.*, 1987). The production of γ-glutamyl hydroxamate was measured in a Hidex Sense microplate reader at 530nm. The absorbance of the reaction mixture was corrected against the absorbance of the negative controls. One unit of the enzyme was defined as the amount of enzyme that catalyses the formation of 1 μmol γ-glutamyl hydroxamate (Bender *et al.*, 1977). The assay was performed on two technical replicates of two biological replicates.

The enzyme activity assay for succinate semialdehyde dehydrogenase (SSA-DH) was adapted after Tian *et al.* (2015). Five μL of nodule extract, which corresponded to 100 ng of soluble protein, were added to 100 μL of reaction buffer (50 mM HEPES (pH 8.0), 1 mM 2-mercaptoethanol, 8 mM NADP^+^ or NAD^+^). The absorbance of the extracts at 340 nm was monitored in a Hidex Sense microplate reader for 60 minutes at 27°C to allow background activity to cease. Then SSA (Santa Cruz Biotechnology, Germany) was added to a final concentration of 4 mM and the absorbance was monitored for an additional 120 minutes. In the negative control, no SSA was added. The measurements were corrected against the background absorbance of buffer with added SSA. The SSA-DH assay was performed on three biological replicates and two technical replicates. The assay was conducted on two independent occasions. All enzyme activity assay data were analysed and visualised in RStudio.

Protein modelling was done through the SWISS-Model portal (https://swiss-model.expasy.org/ visited during September 2021 and March 2022). Visualisation of models was done with UCSF Chimera version 1.14 (Huang *et al.*, 2014; Goddard *et al.*, 2007). For phylogenetic analysis, amino acid sequences of *Frankia*, solved crystal structures of 2-oxoglutarate decarboxylase *Mycobacterium smegmatis* (reference A0R2B1) and the 2-oxoglutarate dehydrogenase of *Staphylococcus epidermis* (reference Q5HPC6) were used to run a BLASTP against the NCBI nr database (March 2022). Sequences were taken from different actinobacterial groups. Sequences were aligned using clusalo on Seaview, and the N terminus was trimmed to the predicted first active domain, as sequences varied in length. Re-aligned sequences were then uploaded to the CIPRES science gateway (Miller *et al.*, 2010). Model of substitution was predicted through ModelTest-NG, and a phylogenetic tree was built using RAxML-HPC2 with 100 bootstraps.

## Results and discussion

### Expression levels of nitrogenase genes allowed for direct comparison of three types of nodules

While the analysis at the transcriptional level has its limitations due to post-transcriptional regulation, it has the advantage to separate plant from bacterial transcription in nodules. Most pathways contain a ratelimiting step catalysed by a key enzyme (Rognstad, 1979). The expression level of the corresponding gene can be the main indicator of the overall activity of the pathway.

Nodules of *Alnus glutinosa* induced by *Frankia alni* ACN14a, *Ceanothus thyrsiflorus* induced by *Candidatus* Frankia californiensis Cv1, and *Datisca glomerata* induced by *Candidatus* F. californiensis Dg2, show various anatomical differences (Pawlowski and Demchenko, 2012). It was unclear whether the contribution of nitrogen-fixing *Frankia* mRNA in total nodule mRNA was similar enough to allow a direct comparison of bacterial gene expression levels. The expression levels of the structural nitrogenase genes *nifDHK* were compared and analysed against the bacterial housekeeping gene *infC*, encoding the transcription initiation factor IF-3 (Alloisio *et al.*, 2010; Nguyen *et al.*, 2019). No significant difference could be found for any of the three genes (p>0.05; Supporting Information Figure S1). It was concluded that a direct comparison of the different nodule types was appropriate. The expression of *nifD* showed the least variation between biological replicates in all treatments. It was used, together with *infC*, to normalise all further gene expression data.

### ROS stress levels are higher in nodules of *D. glomerata* than of *C. thyrsiflorus*

Oxygen protection mechanisms for nitrogenase differ in nodules of *D. glomerata* compared to *C. thyrsiflorus.* The expression levels of genes encoding proteins involved in the detoxification of ROS, catalase, the bacterial haemoglobin *trHbO*, and enzymes synthesising mycothiol -the glutathione equivalent of actinobacteria (Newton *et al.*, 1996)-, were examined. This would determine which oxygen protection mechanism leads to the least ROS stress. The results suggested that ROS stress levels are higher in nodules of *D. glomerata* than of *C. thyrsiflorus* (p<0.05; Supporting Information Figure S2). Our results are consistent with previous data regarding the expression of plant ROS stress-related genes in nodules (Salgado *et al.*, 2018).

### *Frankia* cluster-2 exports different assimilated nitrogen sources depending on the host plant

To determine if an assimilated nitrogen source is exported by *Frankia* cluster-2 to its host plant, ammonium assimilation activities were compared between nodules of *D. glomerata* and *C. thyrsiflorus*. The expression levels of the bacterial genes encoding enzymes of the glutamine synthetase/glutamate synthase (GS/GOGAT) pathway were examined. To examine whether arginine is also involved in *Ceanothus* spp., genes encoding enzymes of the arginine biosynthesis pathway were included in the analysis (Figure 1). For comparison, the expression levels of *Frankia* genes of the GS/GOGAT pathway were also examined in nodules of *A. glutinosa*, where ammonia is known to be exported to the host (*Guan *et al.*, 1996*; Alloisio *et al.*, 2010).

**Figure 1.**
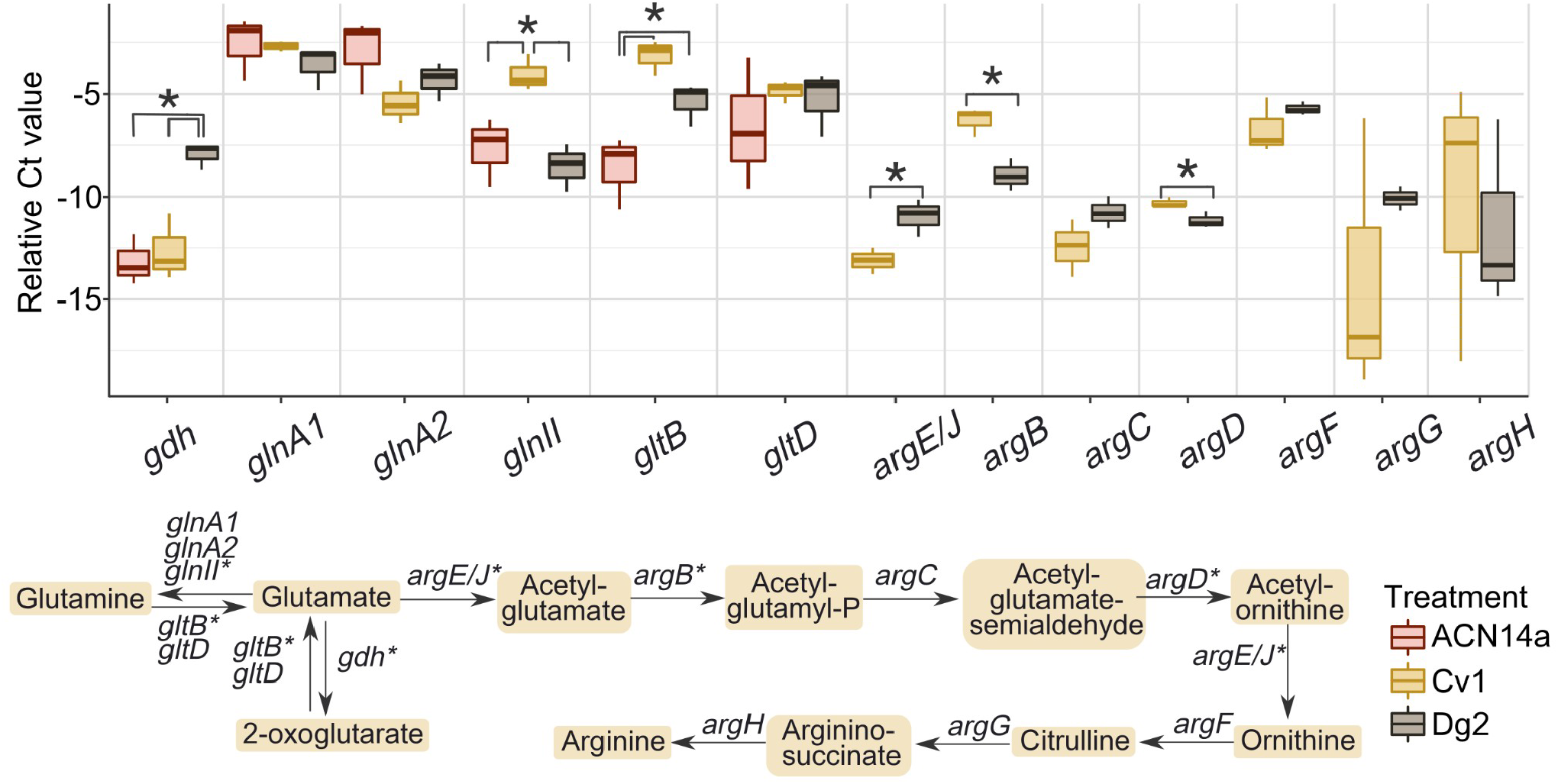
Relative expression levels (delta Ct value) of genes involved in the GS/GOGAT and arginine biosynthesis pathway. The Ct value is normalised against the gene *infC*, encoding the transcription initiation factor IF-3 (Alloisio *et al.*, 2010), and the nitrogenase subunit (MoFe protein) gene *nifD.* An asterisk indicates a significant difference (p <0.5), based one-way ANOVA (ACN14a, Cv1, and Dg2), or on Student’s T-test (Cv1 and Dg2), of gene expression of four technical repeats of three biological repeats of nodules from *Alnus glutinosa* induced by *Frankia alni* ACN14a (red, left), *Ceanothus thyrsiflorus* induced by *Candidatus* Frankia californiensis Cv1 (yellow, centre), and *Datisca glomerata* induced by *Candidatus* Frankia californiensis Dg2 (brown, right). Abbreviations: *gdh*: glutamate dehydrogenase; *glnA1/glnA2*: glutamine synthetase I subunits; *glnII*: glutamine synthetase II; *gltB*: glutamate synthase, large chain; *gltD*: glutamate synthase, small chain; *argB*: acetylglutamate kinase; *argC*: N-acetyl-gamma-glutamyl-phosphate reductase; *argD*: acetylornithine/succinyldiaminopimelate aminotransferase; *argE/argJ*: bifunctional gene acetylornithine deacetylase; *argF*: ornithine carbamoyltransferase; *argG*: argininosuccinate synthase; *argH*: argininosuccinate lyase.

The *gdh* gene, encoding glutamate dehydrogenase which is responsible for the synthesis of 2-oxoglutarate (2-OG) from glutamate, was expressed at significantly higher levels in nodules of *D. glomerata* than of *C. thyrsiflorus* or *A. glutinosa* (p<0.01). *glnA1* and *glnA2* encoding glutamine synthetase I, were found to be expressed at similar levels across all three nodule types (p>0.05). The glutamine synthetase II gene, *glnII*, was expressed at higher levels in nodules of *C. thyrsiflorus* than in nodules of *D. glomerata* (p<0.01) or *A. glutinosa* (p<0.05). The genes encoding the subunits of glutamate synthase, *gltB/gltD*, were not significantly differentially expressed in nodules of *C. thyrsiflorus* and *D. glomerata* (p>0.05). However, *gltB* was expressed at significantly higher levels in both nodules of cluster-2 hosts than in those of *A. glutinosa* (p<0.01 and p<0.05 respectively). Taken together, the results of the expression level of the genes involved in the GS/GOGAT cycle suggest that ammonium assimilation activity is higher in *Frankia* in nodules of *C. thyrsiflorus* and *D. glomerata* than of *A. glutinosa*. This would be consistent with the hypothesis that *Frankia* exports an assimilated source of nitrogen in the former two, but not in the latter.

Looking at the genes encoding enzymes belonging to the arginine biosynthesis pathway, the gene encoding the bifunctional enzyme ArgE/ArgJ was found to be expressed at higher levels in nodules of *D. glomerata* than in *C. thyrsiflorus* (p<0.05; Figure 1). This enzyme catalyses two crucial steps: the synthesis of acetyl-glutamyl-phosphate and the synthesis of ornithine. The first step was shown to be rate-limiting for arginine biosynthesis (Xu *et al.* 2020). Thus, although the genes *argB* and *argD* (p<0.05) were expressed at higher levels in *Frankia* in nodules of *C. thyrsiflorus* than in *D. glomerata*, the results suggest that *Frankia* does not export arginine in nodules of *C. thyrsiflorus*, in contrast with *D. glomerata*. This is consistent with conclusions drawn based on plant gene expression levels (Salgado *et al.*, 2018).

This begs the question: which metabolite is then exported from the bacteria to the plant? To address this question, the levels of different amino acids and nitrogen metabolites were compared between nodules, inoculated roots, and uninoculated roots of *C. thyrsiflorus* (Figure 2). The nitrogen sources that accumulate at the highest levels would represent good candidates for the export by *Frankia* to the host, as previously seen for *D. glomerata* and *A. glutinosa* (Berry *et al.*, 2011; Persson *et al.*, 2016). The results show that in general, the concentrations of nitrogenous metabolites were significantly higher in nodules compared to roots (p<0.05; Figure 2). Consistent with the results on gene expression, and in contrast with the results for nodules of *D. glomerata* (Berry *et al.*, 2004) and *Coriaria myrtifolia* (Coriariaceae, Cucurbitales; Wheeler and Bond, 1970), arginine did not accumulate at high levels. Out of 32 studied metabolites, asparagine and glutamate were the dominant nitrogenous solutes in nodules. These results are in agreement with previously published data on *Ceanothus velutinus* (Wheeler and Bond, 1970).

**Figure 2.**
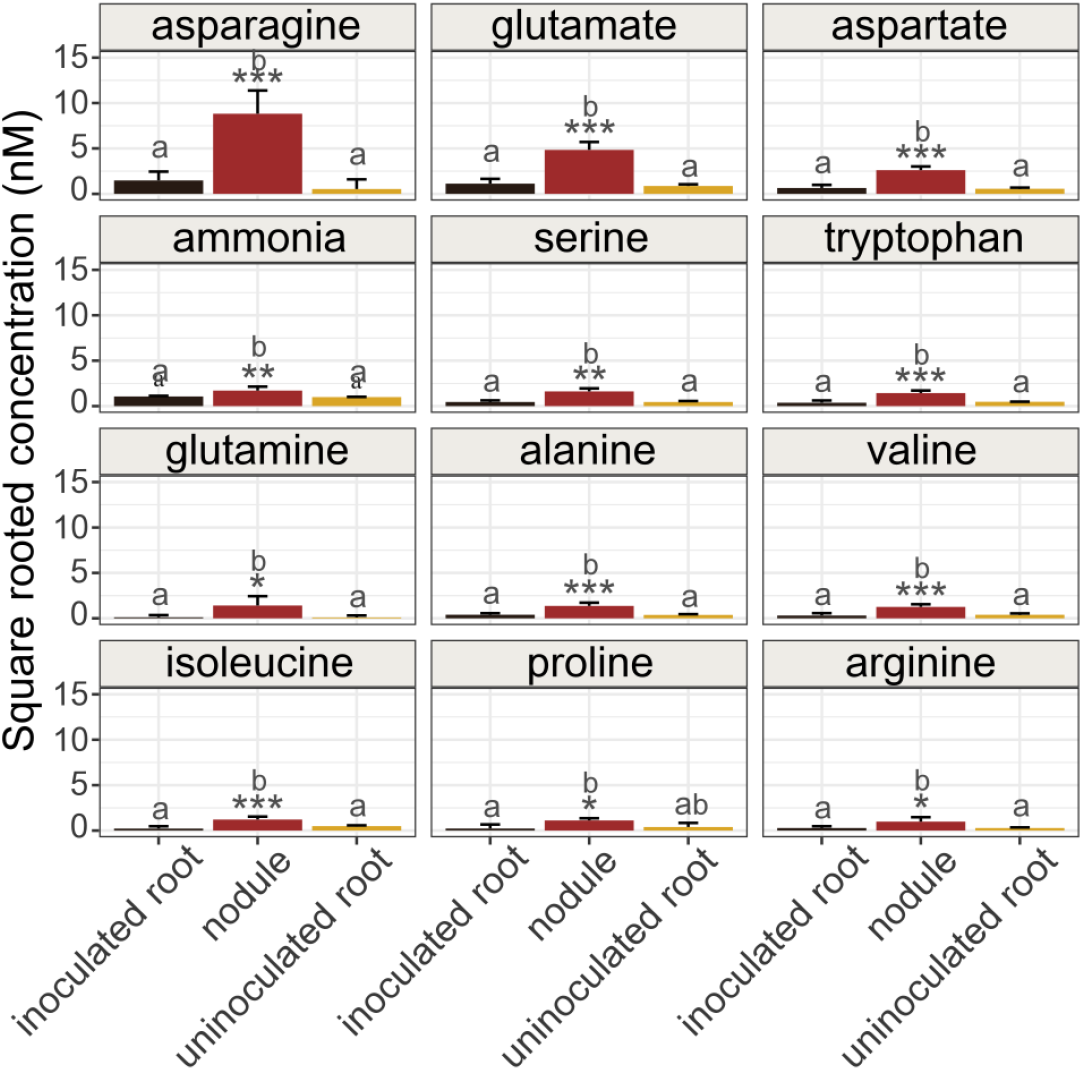
Concentrations of nitrogen metabolites in *Ceanothus thyrsiflorus*. A comparison was made between inoculated roots (dark brown), nodules (red), and uninoculated roots (yellow). The concentration of each metabolite is given as the square root of nM concentration. Only the highest eleven measurements plus the data for arginine are presented. Significant differences are illustrated with stars (* p <0.05; ** p <0.01; *** p <0.005), based on one-way ANOVA of three biological replicates.

The gene encoding glutamine synthetase II (*glnII*) was found significantly higher expressed in nodules of *C. thyrsiflorus* than of *D. glomerata* or *A. glutinosa* (Figure 1). Given the importance of posttranscriptional regulation of GS, an enzyme activity assay was conducted as described previously (Berry *et al.*, 2004), measuring the transferase activity (Figure 3). This would allow determining if ammonium assimilation is taking place at high levels in *Frankia.* Unlike plants, certain prokaryotes such as *Frankia* (Edmans *et al.*, 1987) have been shown to contain two variants of GS: a heat-stable GSI and a heat-labile GSII (Huss-Danell, 1997). It is assumed that GSI and GSII resulted from a gene duplication event that predates the bacteria-eukaryote split (Kumada *et al.*, 1993). Aside from their different heat sensitivity, the two variants also differ in their regulation. Eukaryotes, such as plants, commonly only contain the heat-labile variant of GS. This allows for the distinction between plant and bacterial activity under symbiotic conditions.

**Figure 3.**
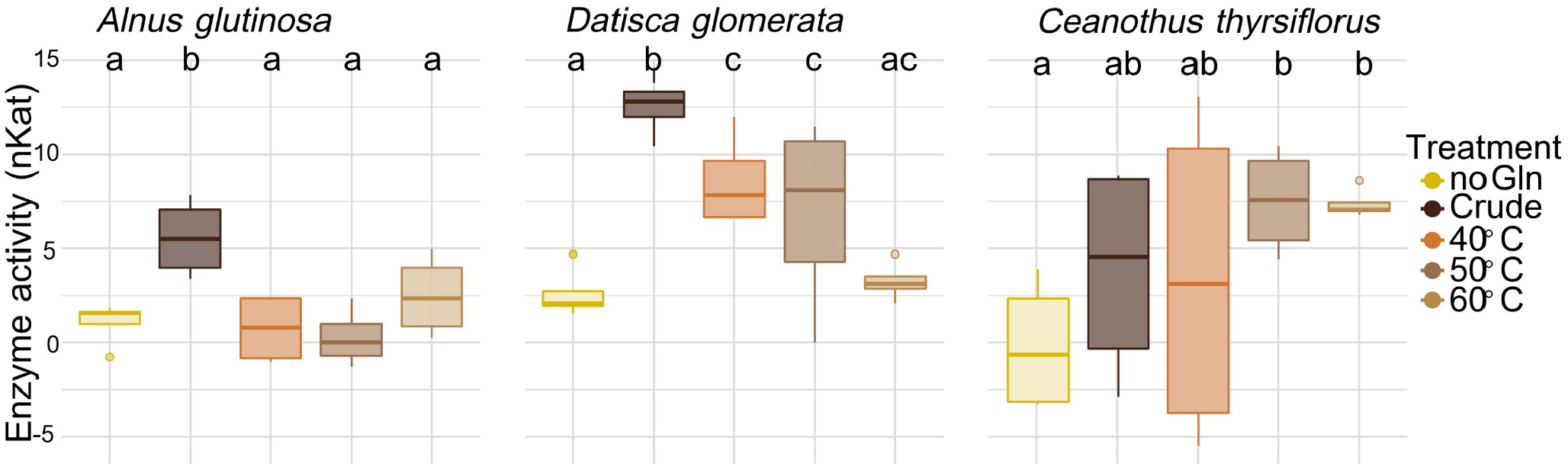
GS activity assay in nodules of *Alnus glutinosa, Datisca glomerata*, and *Ceanothus thyrsiflorus*. *A. glutinosa* was nodulated by *Frankia alni* ACN14a, *D. glomerata* nodulated by *Candidatus* Frankia californiensis Dg2, and *C. thyrsiflorus* by *Candidatus* F. californiensis Cv1. Activity measurements are based on the amount of enzyme required to produce 1μmol of γ-glutamyl hydroxamate (nKat). The activity was measured in either in crude protein extracts, in the negative control without Gln, or in protein extract exposed for 10 minutes to 40°C, 50°C, or 60°C to distinguish between *Frankia* GSI and plant *GS/Frankia* GSII activity. The absorbance at 530nm was corrected against the background absorbance of total denatured protein (boiled for 10 minutes at 95°C). The assay was conducted on two technical replicates of two biological replicates. Letters indicate significant differences (p<0.05, based on one-way ANOVA).

GS activities in nodules of *C. thyrsiflorus* were compared to those in nodules of *A. glutinosa*, where ammonium is exported by *Frankia*, and to *D. glomerata*, where an assimilated source of nitrogen is exported. The heat treatments significantly reduced the GS activity in both *A. glutinosa* and *D. glomerata* (p<0.05) to levels similar to those of the negative control (Figure 3, left and centre panel). This indicates that all measured GS activity in the crude extracts of these plants came from plant GSII, bacterial GSII, or a combination of both, while bacterial GSI activity was below the detection limit. For *A. glutinosa*, this is consistent with previous studies, which indicated that *Frankia* GS activity could not be measured in nodules (Blom *et al.*, 1981), while plant GS was transcribed at high levels in the infected cells (Guan *et al.*, 1996). The results are also consistent with the fact that both plant GS and *Frankia* GS are active in nodules of *D. glomerata* (Berry *et al.*, 2004) while indicating that on the *Frankia* side, GSII plays the determining role. In *C. thyrsiflorus* nodules, however, neither heat treatment significantly reduced the activity of GS (Figure 3, right panel). This indicates that in nodules of *C. thyrsiflorus*, *Frankia* GSI seems to be responsible for ammonium assimilation by *Frankia*, whereas activities of *Frankia* GSII and plant GS are below the detection limit. The undetectable levels of plant GS activity in nodules of *C. thyrsiflorus* support the conclusion that *Frankia* GSI activity is elevated in nodules and *Frankia* exports an assimilated form of nitrogen. Biosynthesis of both asparagine and aspartate relies on a high input of oxaloacetate. Gene expression level of phosphoenolpyruvate carboxykinase (*pepck*) was not elevated in nodules of *C. thyrsiflorus* compared to *D. glomerata* (Figure 4 and Supporting Information Figure S3). However, *Frankia* malate dehydrogenase (*mhd*) expression was significantly elevated in nodules of *C. thyrsiflorus* compared to *D. glomerata* (p=0.01, Figure 4). This could indicate a higher production of oxaloacetate in *Frankia* in nodules of *C. thyrsiflorus* to support the biosynthesis of asparagine and aspartate. The amino acid interconversions between glutamine-glutamate and asparagine-aspartate can occur rapidly and can play important roles in the central metabolism. Analysis of the full transcriptome could provide some more insights. However, the biosynthesis and exchange of asparagine/aspartate and glutamine/glutamate would have to be analysed through enzyme activity assays of *Frankia*, which cannot be achieved under symbiotic conditions, as a distinction from the plant activity cannot be made. Altogether, our results suggest that asparagine and/or glutamate could be exported by *Frankia* cluster-2 in nodules of *C. thyrsiflorus*, and by inference in nodules of other *Ceanothus* spp. They might also play a role as nitrogen transport forms in the xylem. Our data indicate that the export of assimilated nitrogen is a common feature for *Frankia* cluster-2 symbiosis, but which metabolite is exported depends on the host plant order.

**Figure 4.**
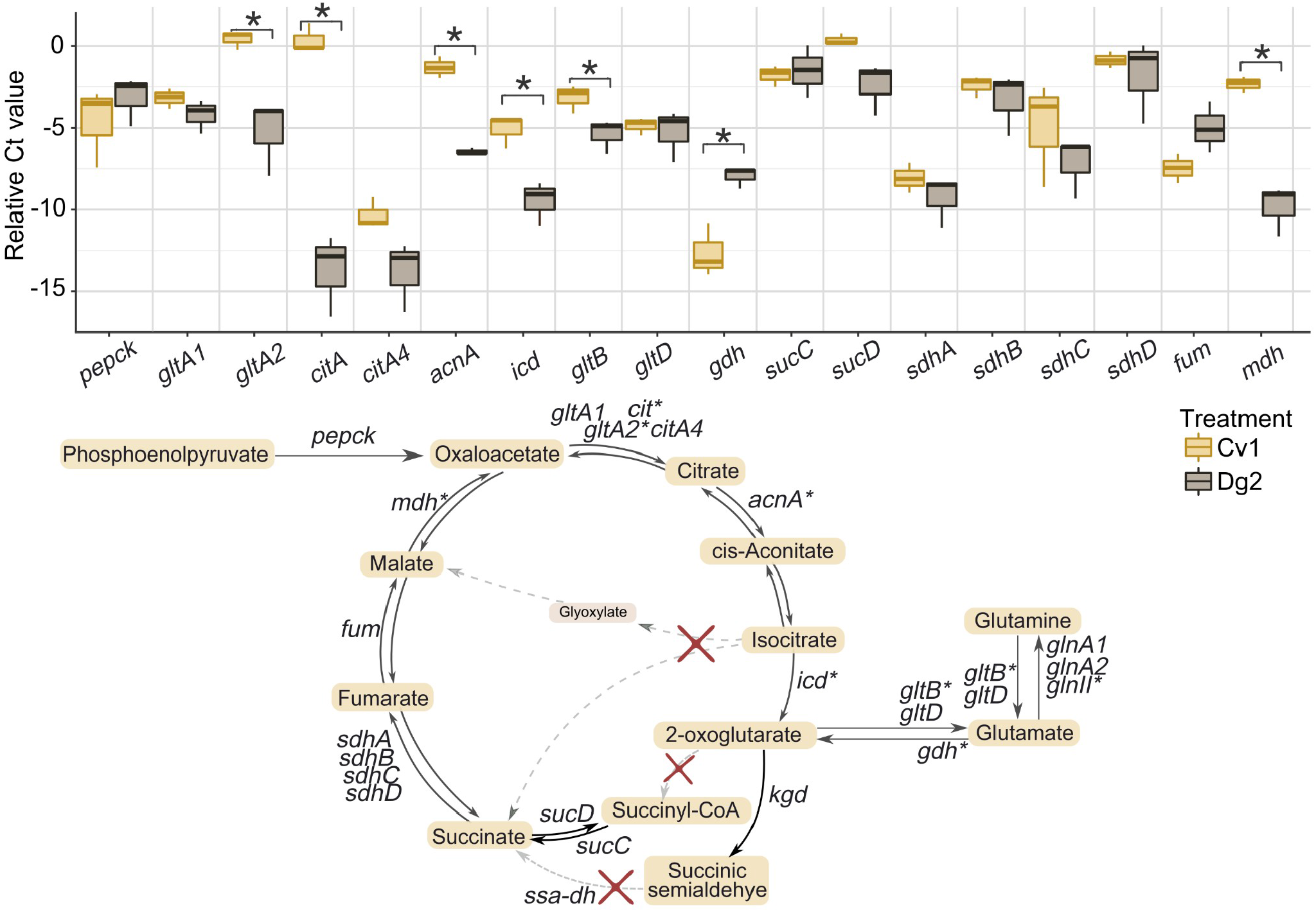
Relative gene expression (delta Ct value) of genes involved in the TCA cycle. The Ct value is normalised against the gene *infC*, encoding the transcription initiation factor IF-3 (Alloisio et al., 2010), and the nitrogenase subunit (MoFe protein) gene *nifD.* An asterisk indicates a significant difference (p <0.5), based on Student’s T-test (Cv1 and Dg2), of gene expression of four technical repeats of three biological repeats of nodules from *Ceanothus thyrsiflorus* induced by *Candidatus* Frankia californiensis Cv1 (yellow, left), and *Datisca glomerata* induced by *Candidatus* F. californiensis Dg2 (brown, right). The pathway with enzymes interacting is illustrated below. Abbreviations used: *pepck*: phosphoenolpyruvate carboxykinase; *gltA/gltA2*: citrate synthase; *citA/citA4*: citrate synthase; *acnA*: aconitate hydratase A; *icd*: isocitrate dehydrogenase; *gltB*: glutamate synthase, large chain; *gltD*: glutamate synthase, small chain; *gdh*: glutamate dehydrogenase; *sucC/sucD*: succinate-CoA ligase subunit alpha/beta; *sdhA/shdB/sdhC/sdhD*: succinate dehydrogenase complex subunit A/B/C/D; *fum*: fumarate hydratase; *mdh*: malate dehydrogenase.

### Gene loss of isocitrate lyase indicates lack of the glyoxylate shunt in *Frankia* cluster-2

The export of assimilated nitrogen, whether in the form of arginine or asparagine, would require a high input of carbon skeletons. Carboxylates provided by the host plant can be used to provide energy or carbon skeletons for ammonium assimilation, both by being shuttled into the TCA cycle. Genome analyses revealed that several genes of TCA cycle-related enzymes were lacking in *Frankia* cluster-2 (indicated by red crosses in Figure 4). The gene *icl*, encoding isocitrate lyase, responsible for catalysing the first step of the glyoxylate shunt (Figure 4), was lacking in all *Frankia* cluster-2 genomes available, in contrast to the genomes of cultivable cluster-1 and cluster-3 strains sequenced to date with the exception of *Frankia casuarinae*. A BLAST search using the sequence of *icl* from *F. alni* ACN14a (Genbank reference: CAJ61004.1), led to the identification of methylisocitrate lyase as the closest homologue in *Candidatus* F. datiscae and in *F. coriariae* (Supporting Information Figure S4), which cannot catalyse the same reaction (Dunn *et al.* 2009). For neither *Candidatus* F. californiensis, nor *Candidatus* Frankia meridionalis, a significant hit could be found. This means that the production of isocitrate will inevitably lead to the synthesis of 2-oxoglutarate (2-OG). Intriguingly, the *icl* gene also seemed to be missing in cluster-1 strains that could never be cultured, such as *Candidatus* Frankia nodulisporulans or *Candidatus* Frankia alpina (Pozzi *et al.*, 2020; Herrera-Belaroussi *et al.*, 2020). Thus, the lack of *icl* could be a common feature for those *Frankia* strains which only have a short phase of saprotrophic growth, and mostly rely on a host plant to support their growth.

### All *Frankia* strains exhibit a variant TCA cycle

2-OG can have different fates as metabolite and signalling molecule, as reviewed by Huergo and Dixon (2015). Within the TCA cycle, 2-OG will be converted to succinate. This can happen by means of three pathways, as explained before: in the ‘classic’ TCA cycle by the activity of 2-OG dehydrogenase, otherwise by the activity of 2-OG decarboxylase (Green *et al.*, 2001; Tian *et al.*, 2005), or through the GABA shunt (Xiong *et al.*, 2014). The latter two pathways require the activity of succinic semialdehyde dehydrogenase (SSA-DH).

The 2-OG dehydrogenase complex is composed of three enzymes: the E1 2-OG dehydrogenase, the E2 dihydrolipoyl succinyltransferase, and the E3 dihydrolipoyl dehydrogenase. Using the corresponding amino acid sequence of *Mycobacteriodes abscessus* and *Staphylococcus epidermis*, the closest homologue of E1 in all *Frankia* genomes examined was found to be a multifunctional 2-OG decarboxylase/oxoglutarate dehydrogenase with less than 45% amino acid sequence identity with the query sequences (accession numbers listed in Supporting Information Table S3). To fully understand if the *Frankia* genes encode E1 dehydrogenase or 2-OG decarboxylase, homology modelling of the amino acid sequence of the *Frankia* proteins was performed on the SWISS Model portal (Figure 5, Supporting Information Figure S5, pdb files available in Supporting information). It was compared to the solved crystal structure of 2-OG decarboxylase from *Mycobacterium smegmatis* (SWISS-MODEL reference A0R2B1) and 2-OG dehydrogenase of *S. epidermis* (SWISS-MODEL reference Q5HPC6). In addition, a phylogenetic tree was built based on the amino acid sequences, which separated the 2-OG dehydrogenase sequences from the 2-OG decarboxylase sequences (Supporting Information Figure S6).

**Figure 5.**
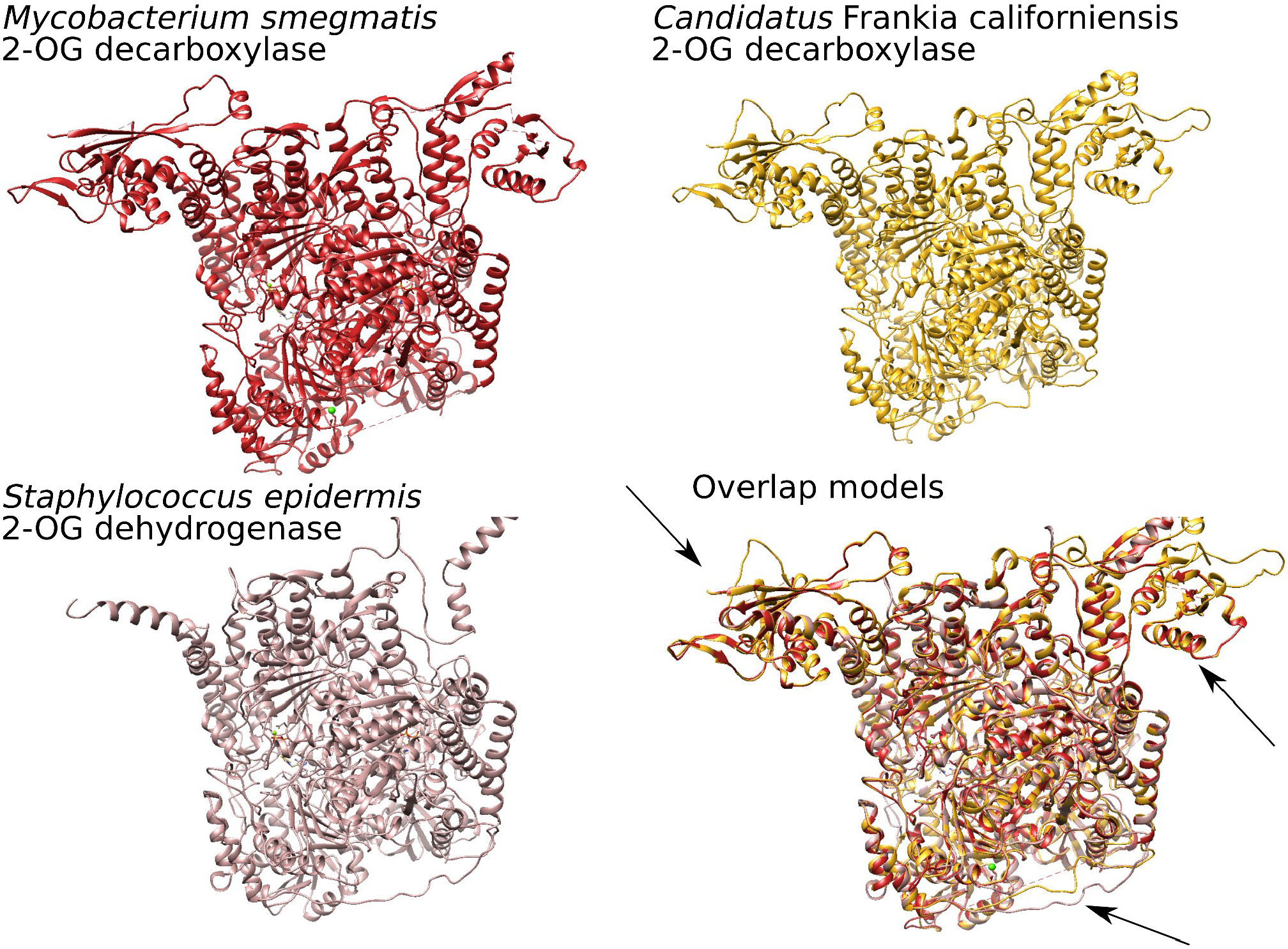
Protein modelling of 2-oxoglutarate decarboxylase. Model of *Candidatus* Frankia californiensis Dg2 (yellow, global model quality estimate QMEANDisCo > 0.70) was compared to the solved crystal structure of *Mycobacterium smegmatis* (red, reference A0R2B1) and to the solved crystal structure of 2-oxoglutarate dehydrogenase of *Staphylococcus epidermis* (rosy brown, reference Q5HPC6). Arrows in the overlay figure highlight the major differences.

The best homology model for all the *Frankia* models was found to be 2-OG decarboxylase (global model quality estimate QMEANDisCo>0.70) while it showed much less similarity to 2-OG dehydrogenase (QMEANDisCo<0.50). The model was highly conserved for all *Frankia* genomes examined, which included all declared species of the different clusters (Supporting Information Table S3, Supporting Information Figure S5, supporting Information pdb files). Furthermore, the model of *Candidatus* F. californiensis Dg2 was compared to solved crystal structure of 2-OG dehydrogenase from *S. epidermis* (Figure 5). While there was overlap, some differences were found, as indicated by arrows in Figure 5. Taken together, we conclude that all *Frankia* genomes available contain a 2-OG decarboxylase gene instead of a 2-OG dehydrogenase. This would imply that the variant TCA cycle is active in which succinate is synthesised from 2-OG via SSA (Figure 4), as it has been shown for some other bacteria. Interestingly, this pathway has been suggested to be distributed widely among rhizobia as an adaptation to the microaerobic conditions for catabolizing dicarboxylic acids (Green *et al.*, 2000). We suggest this adaptation to have evolved in *Frankia* as well.

The production of succinate is thus dependent on the activity of SSA-DH. However, the gene encoding this enzyme could not be identified in *Frankia* cluster-2 genomes. The NADP^+^-dependent variant was found to be common in genomes of strains from other *Frankia* clusters (sequence identity more than 95%, Supporting Information Table S3), whereas the NAD^+^-dependent variant was found only in some of these genomes. Using a BLASTP search with the corresponding sequence from *F. alni* ACN14a, the closest homolog in cluster-2 genomes was found to be a protein belonging to the aldehyde dehydrogenase family (listed in Supporting Information Table S3). Yet, this dehydrogenase had less than 45% amino acid sequence identity with the NADP^+^-dependent SSA-DH in *Frankia* genomes of other clusters. The SSA-DH could also not be identified in the proteome data available for *F. coriariae* BMG5.1 (Ktari *et al.*, 2017).

The lack of strong homologs of NADP^+^-dependent or NAD^+^-dependent SSA-DH alone cannot eliminate the possibility for another enzyme to catalyse the reaction. To determine if any activity was present in *Frankia* cluster-2 strains, an SSA-DH activity assay as described by Tian *et al.* (2005) was conducted for *F. coriariae* BMG5.1, the type strain of the only *Frankia* cluster-2 species able to be grown in culture thus far. *F. alni* ACN14a was used as a positive control because its genome contains genes encoding NADP^+^-dependent as well as NAD^+^-dependent SSA-DH (Supporting Information Table S3). The enzyme activity was determined based on the increase of absorbance due to the production of NADPH or NADH, after adding SSA to the crude protein extract in enzyme buffer. The absorbance significantly increased after the addition of SSA in *F. alni* ACN14a (Figure 6, black solid line after 60 min) in a buffer containing NADP^+^, indicating that the NADP^+^ variant was active. However, a successful assay could not be established for the NAD^+^ variant. In extracts of *F. coriariae* BMG5.1, no activity could be measured after the addition of SSA, either in the presence of NADP^+^ or in the presence of NAD^+^, thus confirming the lack of SSA-DH activity (Figure 6, red). *F. coriariae* BMG5.1 grows considerably slower than any other cultivable *Frankia* strain. The medium recommended by the German Collection of Microorganisms and Cell Cultures (DSMZ, medium 1589; Supporting Information Table S1), contains sodium succinate. This indicates that external supplementation with an intermediate from the reductive part of the TCA cycle is required for this strain to show sufficient growth. The strain was maintained in medium without external succinate for at least three weeks before the activity assay to ensure that SSA-DH activity was not downregulated.

**Figure 6.**
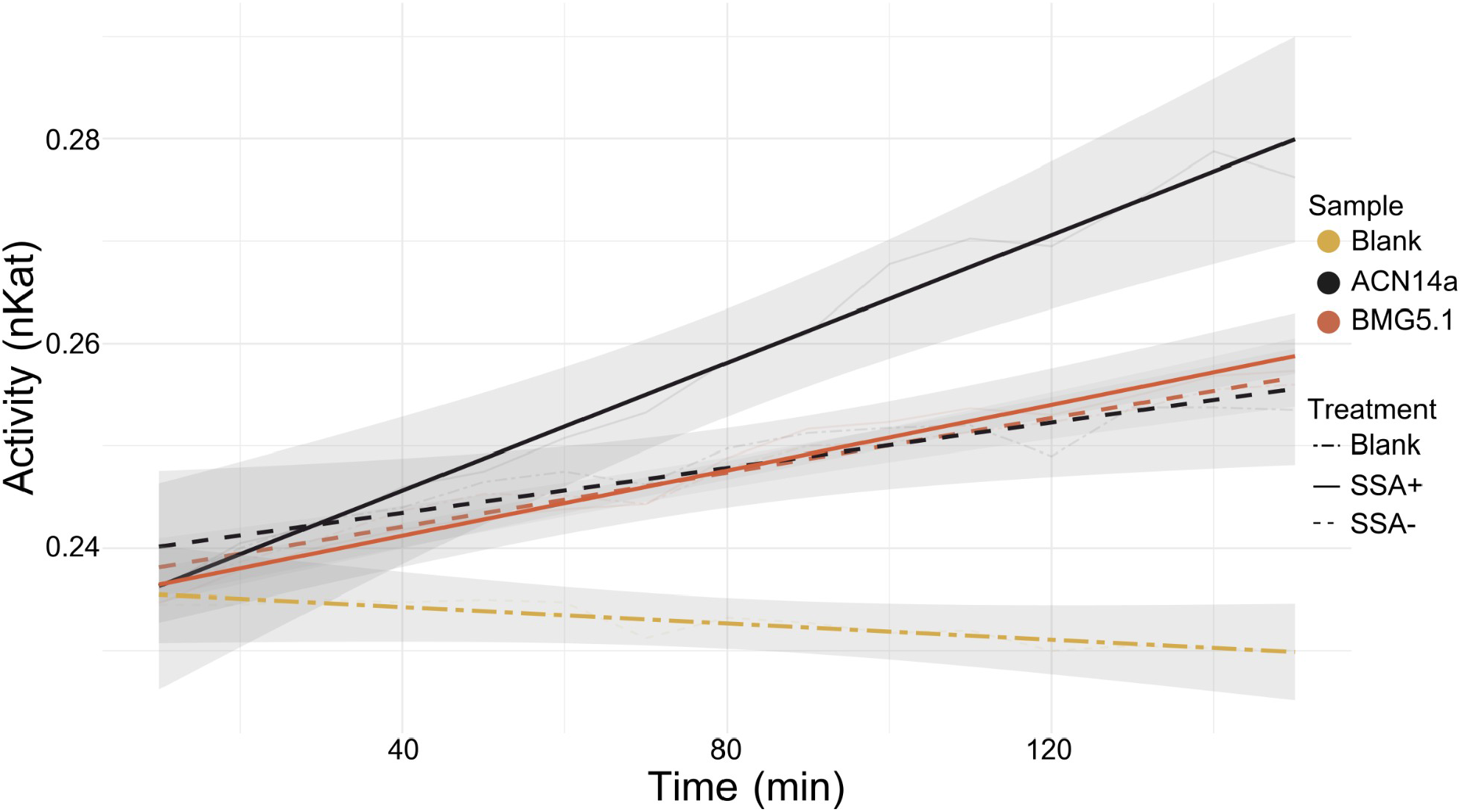
Enzyme activity assay for succinic semialdehyde dehydrogenase (SSA-DH) in *Frankia alni* ACN14a (cluster-1 strain, black) compared to *Frankia coriariae* BMG5.1 (cluster-2 strain, red). Absorbance was measured at 340 nm, to detect the production of NADPH from NADP^+^. Protein extracts were allowed to acclimate for 60 minutes in the reaction buffer with NADP^+^ but without substrate, after which SSA was injected (solid line). Dashed lines indicate negative control, without the addition of SSA, and blank treatment without protein extract (yellow). Grey surface area depicts the standard deviation of three biological replicates and two technical replicates.

Aside from acting as a precursor of succinate, 2-OG can be used for the synthesis of glutamate via the GS/GOGAT cycle, and support ammonium assimilation. As shown above, the export of an assimilated form of nitrogen from the host seems to be a common feature in nodules induced by *Frankia* cluster-2 strains. This would explain the loss of the genes for ICL and SSA-DH observed in *Frankia* cluster-2, as in symbiosis ammonium assimilation would require the majority of 2-OG to be used for ammonium assimilation leading to amino acid biosynthesis. Most reactions of the TCA cycle are reversible, except for the step catalysed by isocitrate dehydrogenase (ICD). The carboxylate provided by the plant as carbon source, which thus far has only been identified for *A. glutinosa* where it represents malate (Jeong *et al.*, 2004), could be converted into any compound on the reductive side of the TCA cycle (Figure 4), thus into fumarate and succinate by the reverse reactions of succinic dehydrogenase; and into fumarate, as well as into oxaloacetate, by malate dehydrogenase. Under symbiotic conditions in nodules, the high input of presumably malate to the reductive side of the TCA cycle would allow 2-OG to be drawn out of the oxidative side of the TCA cycle at a high rate. Hence, the TCA cycle would not be required to work as a cycle, but only as a linear pathway.

We suggest that *Frankia* cluster-2 strains have a unique carbon metabolism: the TCA cycle acts as a linear pathway instead of a cycle to keep up with the constant removal of 2-OG for ammonium assimilation. Gene losses within the TCA cycle are not unique to *Frankia*, as shown by the symbiotic cyanobacteria UCYN-A and *Trichormus azollae* (previously *Nostoc azollae*), which even lack the entire TCA cycle (Tripp *et al.*, 2010; Ran *et al.*, 2010). *Frankia* cluster-2 strains represent the earliest divergent symbiotic clade within the *Frankia* genus. Their metabolism might be based on an ancient form of symbiotic metabolite exchange. The export of an assimilated form of nitrogen is not energy efficient: it requires a much larger supply of carbon skeletons from the host to the endosymbiont. In contrast to ammonia, which can leak through the bacterial membrane and be converted to ammonium in the acidic perisymbiont space, assimilated forms of nitrogen would have to be transported across the bacterial membrane at the expense of energy, which is provided by the host.

The divergence between the symbiotic cluster-2 and the non-symbiotic cluster-4 took place early in the evolution of *Frankia*, followed by the divergence of cluster-1 and cluster-3 from cluster-4 (Nguyen *et al.*, 2016; 2019). *Frankia* cluster-1 strains export ammonium to the host, as opposed to assimilated nitrogen (Guan *et al.*, 1996). This indicates that in different symbiotic clusters the preferred export form evolved differently. The *de facto* linear version of the TCA cycle from the provided carbon source to 2-OG might have led to the loss of genes involved in the production of succinate from 2-OG. The loss of the SSA-DH gene in *Frankia* cluster-2 might have prevented the evolution of a more energy-efficient nutrient exchange system. Given their adaptation of the TCA cycle to the metabolite exchange in symbiosis, it is not surprising that cluster-2 stains have such a low saprotrophic potential and are rarely found in the soil in the absence of a host plant (Battenberg *et al.*, 2017; Persson *et al.*, 2015).

## Conclusions

Based on the data presented in this study, we conclude that the export of an assimilated form of nitrogen by *Frankia* cluster-2 strains in symbiosis is a common feature of the clade. In Cucurbitales host plants, such as *Datisca glomerata* and *Coriaria myrtifolia*, this export form seems to be arginine, while in Rosales, such as *Ceanothus thyrsiflorus*, it seems to be asparagine or glutamate. The assimilation of fixed nitrogen for export during symbiosis puts a high demand on 2-oxoglutarate. The TCA cycle, therefore, seems to work linearly from the carbon source(s) provided by the host, to 2-oxoglutarate. Due to this special metabolism, the need for the glyoxylate shunt is obviated, as well as the production of succinate from 2-oxoglutarate. This led to gene losses which might explain the low saprotrophic potential of *Frankia* cluster-2 strains.

## Acknowledgements

We would like to thank Anna Pettersson and Ingela Lundwall (Stockholm University) for taking care of the plants in the greenhouse, and Rachel Foster and Edouard Pesquet (Stockholm University) for helpful discussions. This project was supported by grants from the Swedish Research Council Vetenskapsradet (VR2012-03061 and VR2029-05540 to KP) and the Carl Tryggers foundation (CTS 9:925 to KP). The bioinformatics support by the BMBF-funded project “Bielefeld-Giessen Center for Microbial Bioinformatics-BiGi (Grant Number 031A533)” within the German Network for Bioinformatics Infrastructure (deNBI.de) is gratefully acknowledged.

## Author contributions

TVN, FB, and KP were responsible for conceptualization, methodology, and investigation. TVN, RH, DW, and FB performed the formal analysis. FB did the visualization. TVN and FB wrote the original draft of the manuscript. All co-authors reviewed, edited and approved the final version. Funding was acquired by KP. FB and TVN contributed equally.

## Data availability

Data are available in the Supporting Information.

## Abbreviations used

2-OG: 2-oxoglutarate
*acnA*: aconitate hydratase A
*argB*: acetylglutamate kinase
*argC*: N-acetyl-gamma-glutamyl-phosphate reductase
*argD*: acetylornithine/ succinyldiaminopimelate aminotransferase
*argE/argJ*: bifunctional gene acetylornithine deacetylase
*argF*: ornithine carbamoyltransferase
argG: argininosuccinate synthase
*argH*: argininosuccinate lyase
BAP: basic propionate
BSA: bovine serum albumin
*citA/citA4*: citrate synthase
*fum*: fumarate hydratase
GABA: gamma-aminobutyrate
*gdh*: glutamate dehydrogenase
*glnA1/glnA2*: glutamine synthetase
*glnII*: glutamine synthetase
*gltA/gltA2*: citrate synthase
*gltB*: glutamate synthase, large chain
*gltD*: glutamate synthase, small chain
GS/GOGAT: glutamine synthetase/glutamate synthase
*icd*: isocitrate dehydrogenase
*mdh*: malate dehydrogenase
*pepck*: phosphoenolpyruvate carboxykinase
PMSF: phenylmethylsulfonylfluoride
*sdhA/shdB/sdhC/sdhD*: succinate dehydrogenase complex subunit A/B/C/D
SSA: succinic semialdehyde
SSA-DH: succinic semialdehyde dehydrogenase
*sucC/sucD*: succinate-CoA ligase subunit alpha/beta
succinyl-CoA: succinyl-coenzyme A
TCA: tricarboxylic acid

