## Supplementary Information for "Gene losses resulting in specialized metabolism in the earliest divergent symbiotic *Frankia* clade can be linked to its low saprotrophic capabilities"

### 1589. modified BAP$^+$ (GFS) MEDIUM FOR FRANKIA

| | | |
|---|---|---|
| $KH_2PO_4$ | 0.950 | g |
| $K_2HPO_4$ | 0.600 | g |
| $NH_4Cl$ | 0.270 | g |
| Na-propionate | 0.480 | g |
| Glucose | 0.480 | g |
| Fructose | 0.480 | g |
| Sodium succinate dibasic | 0.480 | g |
| $MgSO_4$ x 7 $H_2O$ | 0.025 | g |
| $CaCl_2$ x 2 $H_2O$ | 0.010 | g |
| Vitamine solution (see below) | 1.000 | ml |
| Chelated iron solution (see below) | 1.000 | ml |
| Trace element solution (see below) | 0.100 | ml |
| Distilled water | 1000.000 | ml |

*Vitamine solution:*

| | | |
|---|---|---|
| Thiamine HCl | 0.010 | g |
| Nicotinic acid | 0.050 | g |
| Pyridoxine HCl | 0.050 | g |
| Distilled water | 100.000 | ml |

*Chelated iron solution:*

| | | |
|---|---|---|
| Citronensäure | 1.000 | g |
| Eisencitrat | 1.000 | g |
| Distilled water | 100.000 | ml |

*Trace element solution:*

| | | |
|---|---|---|
| $CoSO_4$ x 7 $H_2O$ | 0.001 | g |
| $CuSO_4$ x 5 $H_2O$ | 0.080 | g |
| $H_3BO_3$ | 2.860 | g |
| $MnCl_2$ x 4 $H_2O$ | 1.810 | g |
| $Na_2MoO_4$ x 2 $H_2O$ | 0.025 | g |
| $ZnSO_4$ x 7 $H_2O$ | 0.220 | g |
| Distilled water | 100.000 | ml |

The pH of the final solution is adjusted to 6.3.

Note: Please autoclave the chelated iron solution before you add to the medium, so that solves everything!!!!

© 2007 DSMZ GmbH - All rights reserved

**Supplementary Table S2: primer sequences used**

| Gene | Forward | Reverse |
|---|---|---|
| *acnA* | CATACGTATACGATCCGCCG | GGCCGTGATGTTGGCACCG |
| *argB* | CGTGATCAAGTATGGCGG | CCGCCGTGGACGACGAC |
| *argC* | ACGCCGACATCGTCTTTCTCG | GTCGGCGAGCCGGTGGT |
| *argD* | TGGGCCACGTGTCCAAC | GCAGAAGAACACCCGGC |
| *argE/argJ* | AACCGGGTGCAGGCCGC | GGGCCCGTGCAGGCGTT |
| *argF* | CGGTGGTGCTGCTGTTC | AGCTGGGTGGTCGCCGC |
| *argG* | GGCCTGGAGGAGATCGC | ACGAGGCGGGATGGGTCG |
| *argH* | TGCGGGATCACGTCCGCCA | CGTGCTGGAGGTGGGTCAT |
| *betA* | TCGGTGGTGGAATGGCCGG | ACATGACGGCGACGGGT |
| *betB* | GCCGCGATCGTGCCGTG | GGCGAGTGCTCGTCGGG |
| *eno* | CCGCCGTGATCGACCGGAT | GCGGACTTGTCCGGCGT |
| *fba* | ACGCCGGGAGCGACGGT | TGGGCGAACTCGGCCAG |
| *fum* | TCGAGCACGACTCCATGGG | GGAGATCCGCTGGCCCG |
| *gap* | GTTCACCAAGGCCGAGGAC | ACGTCGTCGTTCACGCCGAT |
| *gdh* | GTTCATGGAGGCGCTGC | TCGGCGGCGACGACCAG |
| *gdh* ACN14a | CTGCTACACGGCCTTCATCC | GACGACGAGATAGCTGTCCTC |
| *glk* | CTCGCCTGGCGTGACGA | CCGAAGCGGTACTCCCC |
| *glnA1* | CGCTTCTGTGACCTTCCT | AACCCGCGTATCGAGGACC |
| *glnA1* ACN14a | CAGGAAGAACTCGATTTCGGGA | CGGATGTTCTGTGACCTGATCA |
| *glnA2* | GGCGTGCTGAAGTCCGT | TGTCCGCCTCGTGGACCC |
| *glnA2* ACN14a | CAGGAAGAACTCGATTTCGGGA | CGGATGTTCTGTGACCTGATCA |
| *glnII* | CCCGTATCGTTGCCGAC | CAGGTGAACACGGGCCGCA |
| *glnII* ACN14a | GAATACATCTGGATCGACGGCA | GCTCCTTGCCCTTCTTGATGAT |
| *gltA* | ACGGCGAGCTGCCGTAGC | TCCAGGAAGCTGCTCTTG |
| *gltB* | AACTGCTCCACCTGGCCGG | AGTGGAACTGGTAGAAGGCCC |
| *gltB* ACN14a | GATCCACGGGTAGACCTTGTG | CCGACGACATCCAGATCAAGAT |
| *gltD* | CGTGCGCATCACCGACTGG | GCCGTTGTGGCAGAACG |
| *gltD* ACN14a | TGAACTCGGGGAAGTTGTTCG | AACATCATCCCCGAGTGGAAC |
| *icd* | GGCATCACCAACCTGCACGT | GCGAGGGTGTCCGCCTC |
| *infC* | CCGCCGGTCTGCAAGCT | TCCTTGATGACCGTCTGG |
| *infC* ACN14a | TTTCCTTGATGACGGTCTGGAC | GTCTGCAAGCTGATGGACTTTG |
| *katA* | TGGCCCAGTTCAACCGG | GCCTTCGTGTACGGGGT |
| *katG* | AGCTGGCCGGACAACGCGA | GGCGACGTTGCCGGCGA |
| *mdh* | AAGCCGGGCATGAGCCG | ACGACGGCCTCGGGCGA |
| *mshA1* | TTCGTGGGGCGTATCCAGC | GCCGACCACCGCGACGA |
| *mshB* | GACGAGGTGATCGGGAC | GGACGAGCACCTCGCCCAT |
| *mshC* | GCTGCCTGGCCGGGGAT | GGGTGATCCCGCAGACGTA |
| *mshD* | GGACGGTGGAGGATCTCCG | CCACCTTCGTCCAGTGG |
| *nifD* | GGACCTCATCTCCATCAG | CATGACGGTGAAGTTGTCGA |
| *nifD* ACN14a | TGATCTCGTCGTCGAAGAACTC | TGAACTACATCTGCACGACCAT |
| *nifH* | TCCACCACTCAGCAGAACACGA | CTGGGCCTTGGAGTGCAGG |
| *nifH* ACN14a | CGGAGGTGACGATGTAGATCTC | TCATCACCTCCATCACCTACCT |
| *nifK* | CGGTCAACCCGCTGAAG | AAGTACGAGGTGCACCCCTG |
| *nifK* ACN14a | CGAAGTTCTTCTCCCGGTACTC | TCCTTGACCACAACGACATCTT |
| *pepck* | ATTTGGTGGTGTGACGGTTC | GGATGAACGTTCCTTTGTCG |
| *pfk1* | GGCATCCCGGTCGTCGG | CAGCCGGTCGATGGCCTC |
| *pgi* | GACCTCGGCCCGGCCAT | GTCGCCTCCCAGATGTC |
| *pgk* | GCTCGCGCAGCTGCTCG | CTCACCGTCGCCCAGCC |
| *pgm* | GATCGGCGTCGTGGAG | GTTGGAGCGGCGCA |
| *pyk* | CCTTCTCCGGGCTGACCTC | ACCGTGGTGTGGACGTC |
| *sdhA* | CGGTCATCGTGGGTGCCG | CCGGTGTGGGAGCGGGT |
| *sdhB* | TGCGCGCACGGCGTGTG | CTTGATGGGTTCGATCGAGATTT |
| *sdhC* | CTGTTCATCATGAACATTCTTGA | CGAGCATGAGCAGGTCC |
| *sdhD* | GACACCGTGATCGCGAC | TCCGGATCCCGTTGAGC |
| *sucC* | AAGACCGGCGCCGACGT | ACGCCCTCGGTGATGACCA |
| *sucD* | CGAAGAAGCTGTTCGCCGAC | GCGCTTTGACGACGACACG |
| *trHbO* | CACCTTCTACGACGCGG | TCCGGGTAGAGCGGGCG |

**Supplementary Table S3: gene accession numbers**

| **Gene** | **2-OG decarboxylase/2-OG dehydrogenase** | **SSA-DH NADP+** | **SSA-DH NAD+** |
|---|---|---|---|
| ***M. abscessus*** | SKR88055.1 | | |
| ***S. epidermis*** | WP_194086144.1 | | |
| ***Ca.* F. datiscae** | WP_131765556.1 | WP_206443133.1 | WP_131766698.1 |
| ***Ca.* F. californiensis** | SBW19759.1 | SBW22860.1 | SBW22860.1 |
| ***Ca.* F. meridionalis** | WP_131745573.1 | WP_131745719.1 | WP_131745719.1 |
| ***F. coriariae*** | WP_047222684.1 | WP_047223263.1 | WP_047223244.1 |
| ***F. alni*** | CAJ64606.1 | CAJ64985.1 | WP_083867058.1 |
| ***F. torreyi*** | WP_044885360.1 | | |
| **F. inefficax** | WP_013422021.1 | | |
| ***Ca.* F. nodulisporulans** | WP_163547121.1 | | |
| ***F. saprophytica*** | WP_007518568.1 | | |
| ***F. asymbiotica*** | WP_076812160.1 | | |
| ***F. elaeagni*** | WP_026310503.1 | | |
| ***F. discariae*** | WP_026239651.1 | | |
| ***F. casuarinae*** | OAA21696.1 | | |
| ***F. canadensis*** | WP_101829637.1 | | |

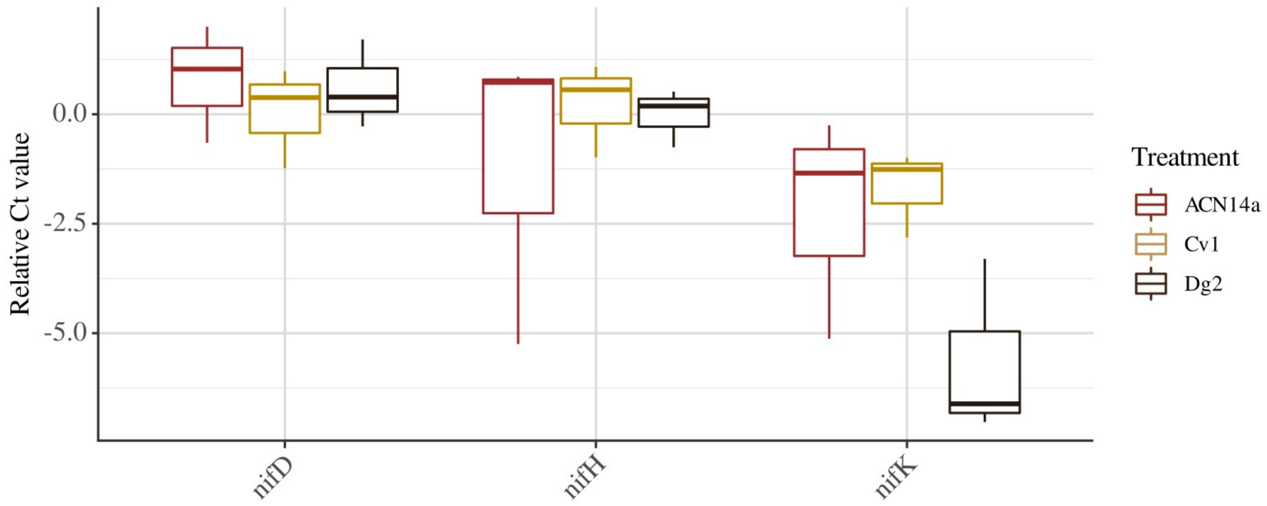


**Supplementary Figure S1: Relative expression levels (delta Ct value) of *nif* genes encodiging the nitrogenase enzyme complex.** The Ct value is normalised against the gene *infC*, encoding the transcription initiation factor IF-3 (Alloisio *et al.*, 2010). An asterisk indicates a significant difference (p <0.5), based one-way ANOVA of gene expression of four technical repeats of three biological repeats of nodules from *Alnus glutinosa* induced by *Frankia alni* ACN14a (red, left), *Ceanothus thyrsiflorus* induced by *Candidatus* Frankia californiensis Cv1 (yellow, centre), and *Datisca glomerata* induced by *Candidatus* Frankia californiensis Dg2 (brown, right).

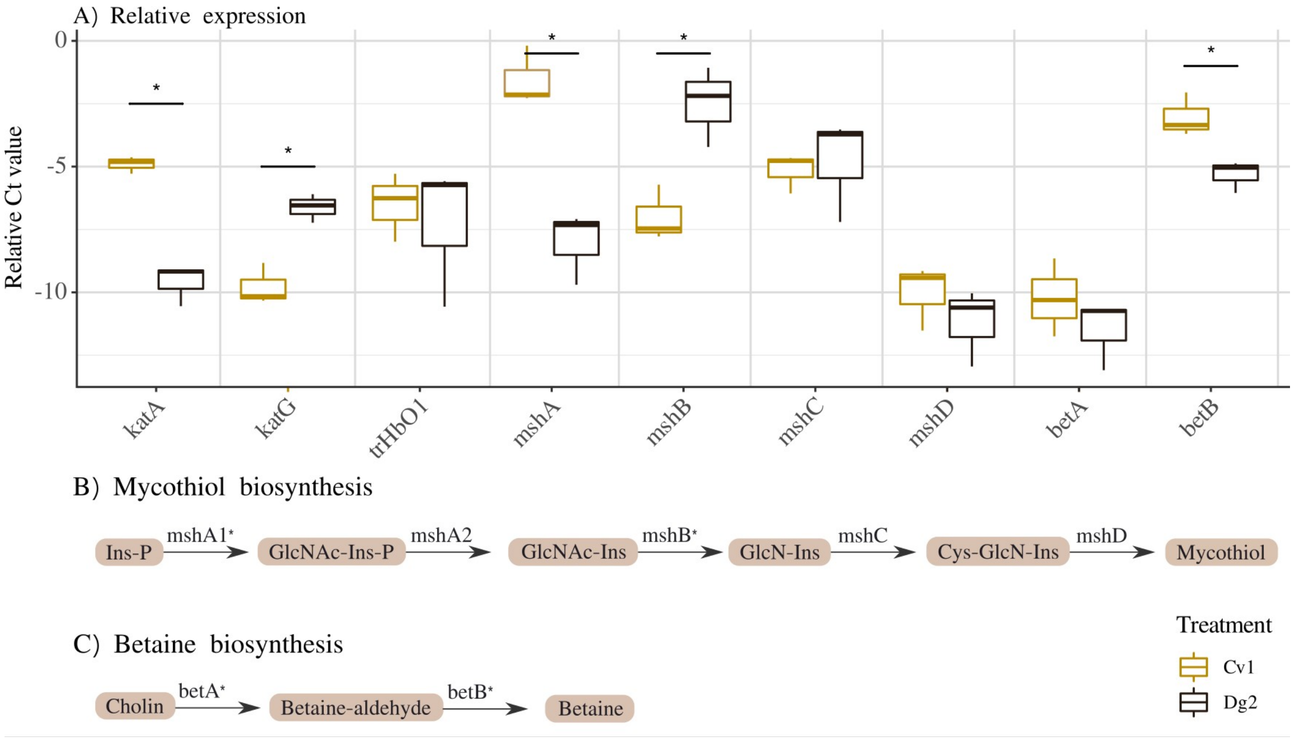


**Supplementary Figure S2: Relative expression levels (delta Ct value) of katalase, hemoglobin, and mycothiol biosynthesis genes encodiging the nitrogenase enzyme complex.** The Ct value is normalised against the gene *infC*, encoding the transcription initiation factor IF-3 (Alloisio *et al.,* 2010), and the nitrogenase subunit (MoFe protein) gene *nifD*. An asterisk indicates a significant difference (p <0.5), based Student's T-test of gene expression of four technical repeats of three biological repeats of nodules from *Ceanothus thyrsiflorus* induced by *Candidatus* Frankia californiensis Cv1 (yellow, left), and *Datisca glomerata* induced by *Candidatus* Frankia californiensis Dg2 (brown, right).

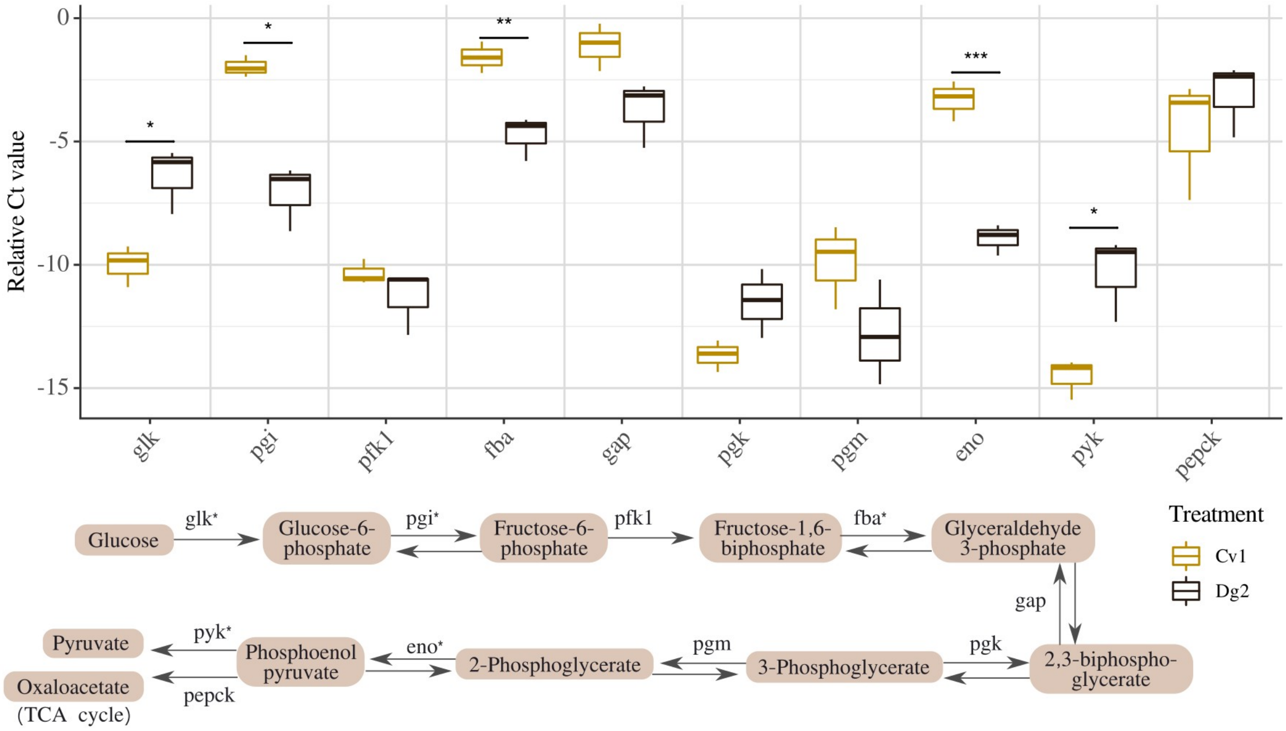


**Supplementary Figure S3: Relative expression levels (delta Ct value) of glycolysis genes.** The Ct value is normalised against the gene *infC*, encoding the transcription initiation factor IF-3 (Alloisio *et al.,* 2010), and the nitrogenase subunit (MoFe protein) gene *nifD*. An asterisk indicates a significant difference (p <0.5), based Student's T-test of gene expression of four technical repeats of three biological repeats of nodules from *Ceanothus thyrsiflorus* induced by *Candidatus* Frankia californiensis Cv1 (yellow, left), and *Datisca glomerata* induced by *Candidatus* Frankia californiensis Dg2 (brown, right).

```
Seaview [blocks=10 fontsize=10 A4] on Mon Sep  6 14:56:15 2021

                      1
F._alni_ACN14a        QAAEELAARW ATDPRWAGIE RTYGAAEVIK LRGSVLEEHT LARRGAEKLW GLLHDTDYVH
Ca_F._datiscae        ---------- ---------- ---------- ----MLHART PAHARRAALR AALRSGRLLR
F._coriariae_BMG5.1   ---------- ---------- ---------- ----MLHART PAHVRRAALR AALCSGRLLR

                     61
F._alni_ACN14a        ALGALTGNQA VQQVKAGLRA IYLSGWQVAA DANLSGQTYP DQSLYPANSV PAVVRRINNA
Ca_F._datiscae        FPGAFNPLSA VLITELGFDG IYVSGAVLSA DLALPD---- -IGLTTLTEV ----------
F._coriariae_BMG5.1   FPGAFNPLSA VLITELGFDG IYVSGAVLSA DLALPD---- -IGLTTLTEI ----------

                    121
F._alni_ACN14a        LLRADQITWA EGVADAPDWL VPIVADAEAG FGGVLNAFEL MSAMITAGAA GVHWEDQLAS
Ca_F._datiscae        AGRAGQIA-- ------RVSD LPALVDADTG FGEPMNVART IQTLEDTGLA GCHIEDQVN-
F._coriariae_BMG5.1   AGRAGQIA-- ------RVSD LPALVDADTG FGEPMNVART IQILEDAGLA GCHIEDQVN-

                    181
F._alni_ACN14a        EKKCGHLGGK VLIPTGQHIR TLNAARLAAD VADVPSVIVA RTDAQAATLI TSDVDE----
Ca_F._datiscae        PKRCGHLDGK TVVPVEEMVR RIRAAVTARR --DENFLICA RTDARAVEGL GGAADRARAY
F._coriariae_BMG5.1   PKRCGHLDGK TVVPVEEMVR RIRAAVTARR --DENFLICA RTDARAVEGL SGAADRARAY

                    241
F._alni_ACN14a        ----RDLPFV TGERTAEGFY RVRDGVEPCI ARGLAYAPYA DLLWMETSTP DLEIARQFAE
Ca_F._datiscae        ADAGADMIFP EAMADAGEFE TVRRAVDVPI LANMTEFGKS ELLTTDT--- -LES----AG
F._coriariae_BMG5.1   ADAGADMIFP EAMADAGEFE TVRRAVDVPI LANMTEFGKS ELLTTDT--- -LES----AG

                    301
F._alni_ACN14a        AIKAQYPDQM LAYNCSP-SF NWRAHLDDAT IAKFQRELGH MGYKFQFITL AGFHALNHSM
Ca_F._datiscae        VSVVIYPVTL LRLAMGAVED GLRRILADGT QAGLVDRMQT RARLYELLDY PGYNTFDTNI
F._coriariae_BMG5.1   VSVVIYPVTL LRLAMGAVED GLRRIRADGT QAGLVDRMQT RARLYELLDY PGYNAFDTNI

                    361
F._alni_ACN14a        FSLAHGYARE GMTAYVDLQE REFASEADGY TATRHQREVG TGYFDLVSTA INPTSDTVAL
Ca_F._datiscae        FNFRI----- ---------- ---------- ---------- ---------- ----------
F._coriariae_BMG5.1   FNFRI----- ---------- ---------- ---------- ---------- ----------

                    421
F._alni_ACN14a        RGSTEEAQF
Ca_F._datiscae        ---------
F._coriariae_BMG5.1   ---------
```

**Supplementary Figure S4: alignment of isocitrate lyase gene.** Isocitrate lyase was taken from *Frankia alni* ACN14a genome and a blast search was performed on cluster-2 genomes. Aligment with the best hit found for *Candidatus* Frankia datiscae Dg1 and *Frankia coriariae* BMG5.1 is shown. No significant hits could be identified for *Candidatus* Frankia californiensis Dg2 or *Candidatus* Frankia meridionalis Cppng1. Alignment was visualised in Seaview.

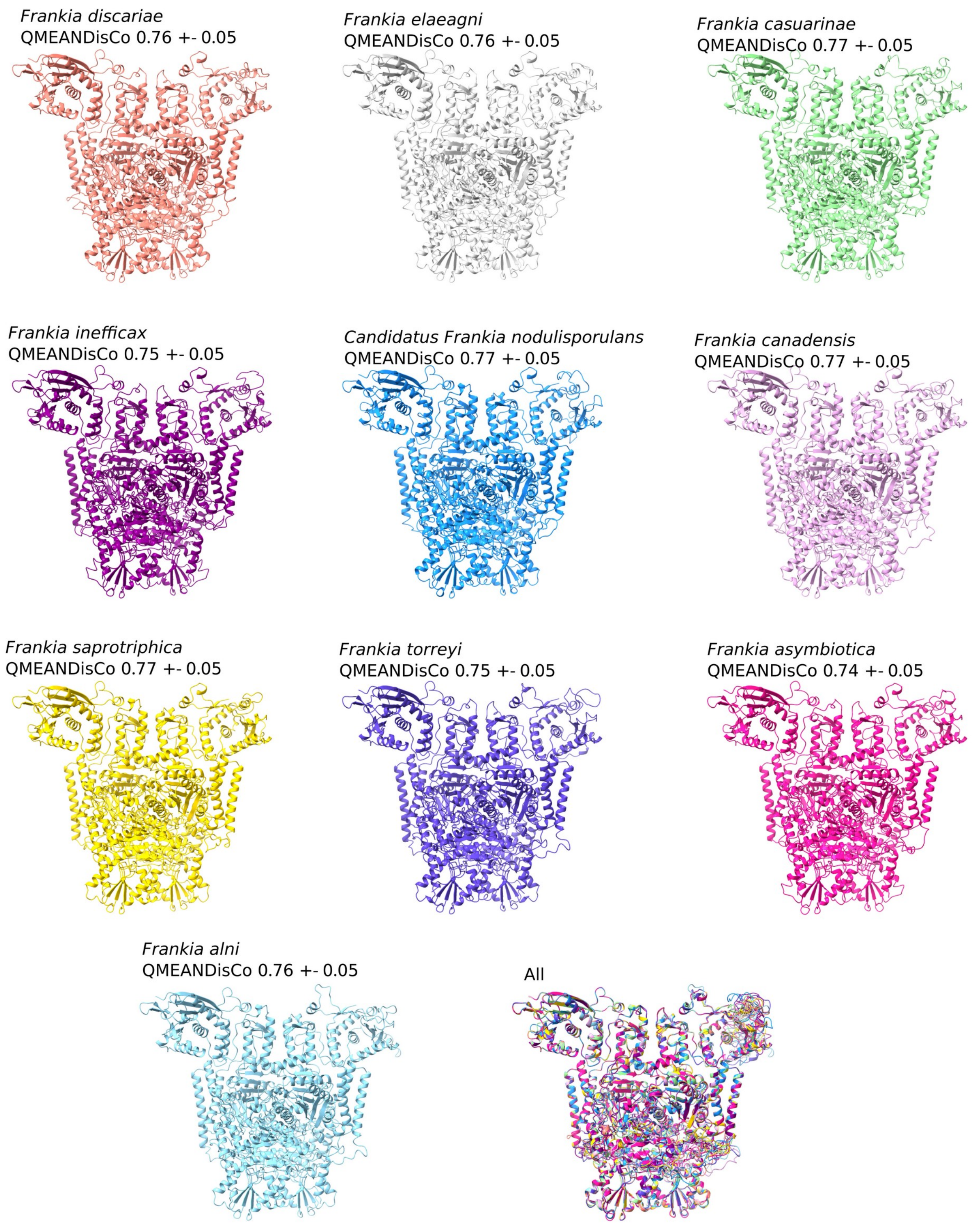


**Supplementary Figure S5: models of 2-oxoglutarate decarboxylase in *Frankia.*** Models were built using SWISS-MODEL portal and visualised in ChimeraX. The global QMEANDisCo for each model is given.

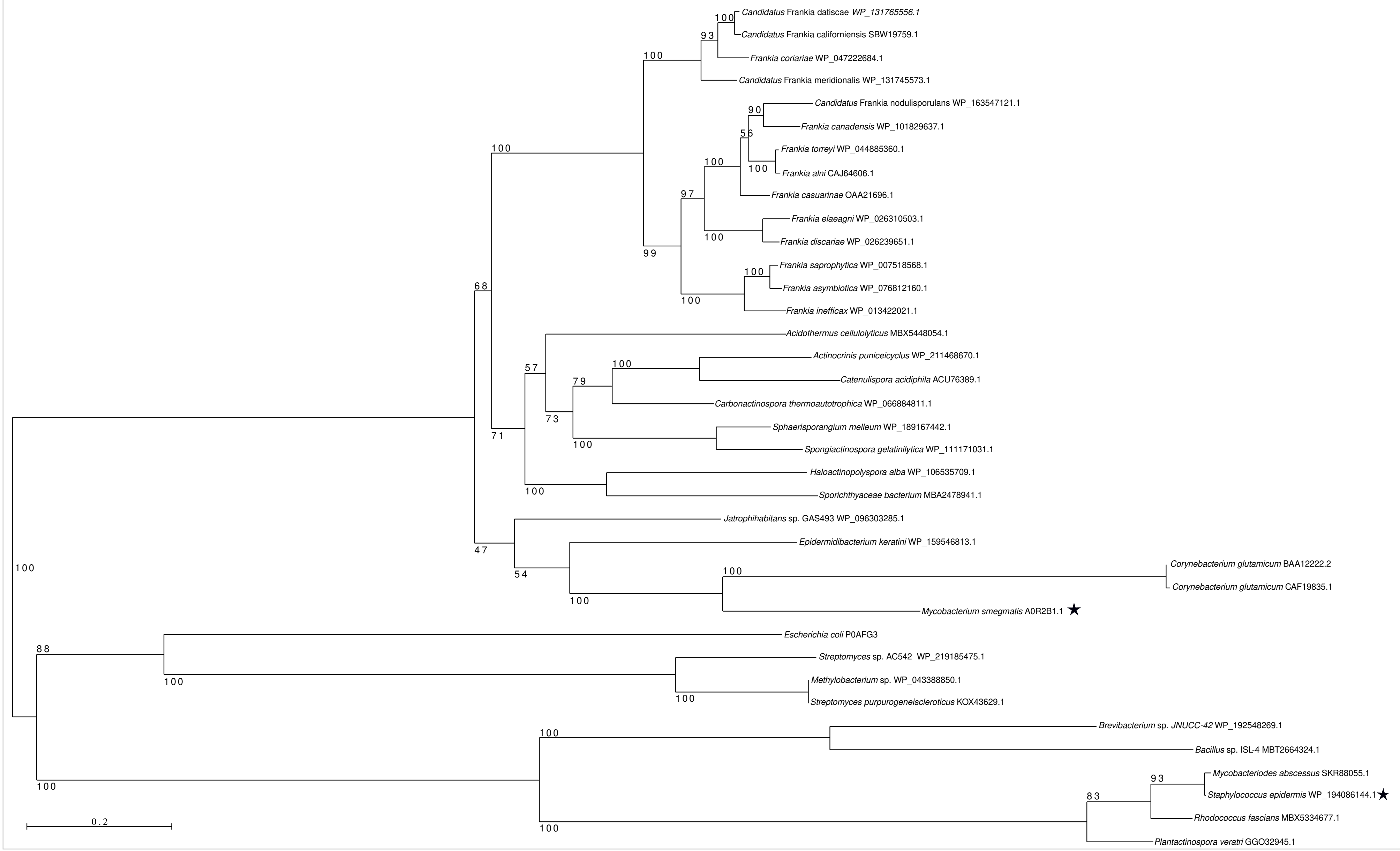

Candidatus Frankia datiscae WP_131765556.1
Candidatus Frankia californiensis SBW19759.1
Frankia coriariae WP_047222684.1
Candidatus Frankia meridionalis WP_131745573.1
Candidatus Frankia nodulisporulans WP_163547121.1
Frankia canadensis WP_101829637.1
Frankia torreyi WP_044885360.1
Frankia alni CAJ64606.1
Frankia casuarinae OAA21696.1
Frankia elaeagni WP_026310503.1
Frankia discariae WP_026239651.1
Frankia saprophytica WP_007518568.1
Frankia asymbiotica WP_076812160.1
Frankia inefficax WP_013422021.1
Acidothermus cellulolyticus MBX5448054.1
Actinocrinis puniceicyclus WP_211468670.1
Catenulispora acidiphila ACU76389.1
Carbonactinospora thermoautotrophica WP_066884811.1
Sphaerisporangium melleum WP_189167442.1
Spongiactinospora gelatinilytica WP_111171031.1
Haloactinopolyspora alba WP_106535709.1
Sporichthyaceae bacterium MBA2478941.1
Jatrophihabitans sp. GAS493 WP_096303285.1
Epidermidibacterium keratini WP_159546813.1
Corynebacterium glutamicum BAA12222.2
Corynebacterium glutamicum CAF19835.1
Mycobacterium smegmatis A0R2B1.1 ★
Escherichia coli P0AFG3
Streptomyces sp. AC542 WP_219185475.1
Methylobacterium sp. WP_043388850.1
Streptomyces purpurogeneiscleroticus KOX43629.1
Brevibacterium sp. JNUCC-42 WP_192548269.1
Bacillus sp. ISL-4 MBT2664324.1
Mycobacteriodes abscessus SKR88055.1
Staphylococcus epidermis WP_194086144.1 ★
Rhodococcus fascians MBX5334677.1
Plantactinospora veratri GGO32945.1
100
93
100
100
90
56
100
100
97
100
99
100
100
68
57
100
79
73
100
71
100
47
54
100
100
100
88
100
100
100
100
93
83
100
0.2

**Supplementary Figue S6: RAxML Maximum likelihood phylogeny for the 2-oxoglutarate dehydrogenase and 2-oxoglutarate decarboxylase.** Sequences were aligned with clustalO in Seaview. Rapid bootstrapping was performed using 100 bootstrap. LG substitution matrix was used. Black asterisk indicates the position of 2-OG decarboxylase from *Mycobacterium smegmatis* (reference A0R2B1) and 2-OG dehydrogenase of *Staphylococcus epidermis* (reference Q5HPC6). Sequence references are given after each name.
